# Olive mild mosaic virus coat protein and p6 are suppressors of RNA silencing and their silencing confers resistance against OMMV

**DOI:** 10.1101/329920

**Authors:** CMR Varanda, P Materatski, MD Campos, MIE Clara, G Nolasco, MR Félix

## Abstract

RNA silencing is an important defense mechanism in plants, yet several plant viruses encode proteins that suppress it. Here the genome of Olive mild mosaic virus (OMMV) was screened for silencing suppressors using a green fluorescent based transient suppression assay. The full OMMV cDNA and 5 different OMMV open reading frames (ORFs) were cloned into Gateway binary destination vector pK7WG2, transformed into *Agrobacterium tumefaciens* C58C1 and agroinfiltrated into *Nicotiana benthamiana* 16C plants. Among all ORFs tested, CP and p6 showed suppressor activity, with CP showing a significant higher activity when compared to p6, yet lower than that of the full OMMV. This suggests that OMMV silencing suppression results from a complementary action of both CP and p6.

Such discovery led to the use of those viral suppressors in the development of OMMV resistant plants through pathogen-derived resistance (PDR) based on RNA silencing. Two hairpin constructs targeting each suppressor were agroinfiltrated in *N. benthamiana* plants which were then inoculated with OMMV RNA. When silencing of both suppressors was achieved, a highly significant reduction in viral accumulation and symptom attenuation was observed as compared to that seen when each construct was used alone, and to the respective controls, thus showing clear effectiveness against OMMV infection. Data here obtained indicate that the use of both OMMV viral suppressors as transgenes is a very efficient and promising approach to obtain plants resistant to OMMV.

**Importance:** OMMV silencing suppressors were determined. Among all ORFs tested, CP and p6 showed suppressor activity, with CP showing a significant higher activity when compared to p6, yet lower than that of the full OMMV, suggesting a complementary action of both CP and p6 in silencing suppression.

This is the first time that a silencing suppressor was found in a necrovirus and that two independent proteins act as silencing suppressors in a member of the *Tombusviridae* family.

When silencing of both suppressors was achieved, a highly significant reduction in viral accumulation and symptom attenuation was observed as compared to that seen when each was used alone, thus showing clear effectiveness against OMMV infection. A high percentage of resistant plants was obtained (60%), indicating that the use of both OMMV viral suppressors as transgenes is a very efficient and promising approach to obtain plants resistant to OMMV.

## Introduction

RNA silencing is a gene inactivation mechanism identified in most eukaryotes that is involved in several biological processes such as regulating endogenous gene expression, maintenance of genome stability and defence against viruses (1, 2).

Amongst the several strategies plants have developed to counter virus infections, RNA silencing is one of the most important (1, 3). In antiviral RNA silencing, double stranded RNAs (dsRNAs) of sizes varying from 30 nucleotides (nt) to hundreds of nt (4) derived from ssRNA virus replication intermediates, are recognized as foreign. Such dsRNAs in plants are first processed by an RNase III-like nuclease (termed DICER-like or DCL) into double stranded viral short interfering RNAs (siRNAs) of 21 to 22 nt long with 2 nt 3’ overhangs (1, 5). Plant virus infections are associated with the accumulation of these virus specific siRNAs. The cleavage is then accomplished by members of the Argonaute protein family (AGOs) (6) which recruit siRNA and associated proteins to form the RNA induced silencing complex (RISC) with one of the siRNA strands (the guide strand) in a process that is accompanied by the release/degradation of the other “passenger” strand.

RISC possesses ribonuclease activity and is guided by the single stranded siRNAs to its target based on sequence complementarity, resulting in binding and degradation of homologous RNA molecules by the catalytic activity of the AGO (7, 8).

Cleavage products of the target RNAs serve as template of RNA dependent RNA polymerases (RDR) to form dsRNAs, leading to secondary siRNA production (9), which can again initiate silencing in a self-sustained manner.

To counteract host RNA silencing defense, viruses have evolved several strategies. One of such involves viral proteins encoded in the genomes that suppress plant RNA silencing, termed viral suppressors of RNA silencing (VSRs) (10, 11). Most viruses studied have one VSR and many VSRs have been identified (12). These proteins are highly divergent, appearing to have evolved independently in the different viruses and they may interfere in different stages of the RNA silencing pathways, either binding dsRNAs and inhibiting its processing into siRNAs; or sequestering viral siRNAs preventing their incorporation into RISC or directly interfering with recognition of viral RNA, dicing and RISC assembly (8, 13).

Viruses from different genera within *Tombusviridae* family, have different suppressors. The p19 of tombusviruses *Tomato bushy stunt virus* (TBSV) and *Cymbidium ringspot virus* (CymRSV), associated with long distance movement, is one of the most studied viral suppressors (14-16). The only function of the p14 of the aureusvirus, such as *Pothos latent virus*, seems to be to prevent silencing (17).

The CP of betacarmoviruses such as *Turnip crinkle virus*, as well as the movement proteins of some dianthoviruses also act as silencing suppressors (18-21). No silencing suppressors are identified in the other genera in the family *Tombusviridae*, which would help to elucidate the evolutionary progression of viruses within this family.

OMMV is a member of the genus *Alphanecrovirus* within the *Tombusviridae* family, being one of the most spread viruses in olive orchards (22-24). Its genomic RNA is 3683 nt long with 5 open reading frames (ORFs) and the virus is likely to have resulted from recombination events between two other necroviruses OLV-1 and TNV-D, based on the high amino acid identity with the RNA dependent RNA polymerase of OLV-1 and of the CP of TNV-D (25). ORF1 (p23), pre-readthrough, and ORF2 (p82), predicted as the RdRp, are involved in RNA replication, ORF3 (p8) and ORF 4 (p6) are predicted to be involved in virus movement, and ORF5 (p29) is predicted as the CP and it is involved in capsid assembly, systemic movement and vector transmission (26). To the best of our knowledge, no other functions of the gene products such as RNA silencing suppression were ever identified in any alpha- or betanecrovirus.

In this study, OMMV encoded proteins were examined to identify suppressors of RNA silencing and two were found, the CP and p6. OMMV silencing suppression ability seems to result from a coordinate and complementary action of both. In addition, resistance to OMMV in *N. benthamiana* was achieved using hpRNA constructs containing both CP and p6.

## Results

### Determination of OMMV silencing suppressors

To identify potential OMMV RNA silencing suppressors, all proteins encoded by the genome were tested for their ability to suppress silencing by using a GFP reporter gene in plant tissues. Individual OMMV ORFs were cloned into a binary vector (pk7wg2) driven by the CaMV 35S promoter in *A. tumefaciens*. For comparison, a full-length cDNA clone of OMMV that generates a full infection in *N. benthamiana* was also cloned. Transient expression assays were performed on transgenic 16C *N. benthamiana* plants by co-infiltrations of each clone plus pK_GFP, a construct that expresses a transcript homologous to the transgene of 16C plants.

Infiltrated *N. benthamiana* leaf patches showed GFP fluorescence under UV light at 2 dpi in all samples (data not shown). Leaves infiltrated with the silencing inducer pk_GFP showed a weak GFP fluorescence at 3 dpi (Figure 1, GFP 3dpi) and at 5 dpi fluorescence disappeared (Figure 1, GFP 5 dpi) and silencing began, as seen by the development of a red colour under UV light on the infiltrated region.

**Figure 1.**
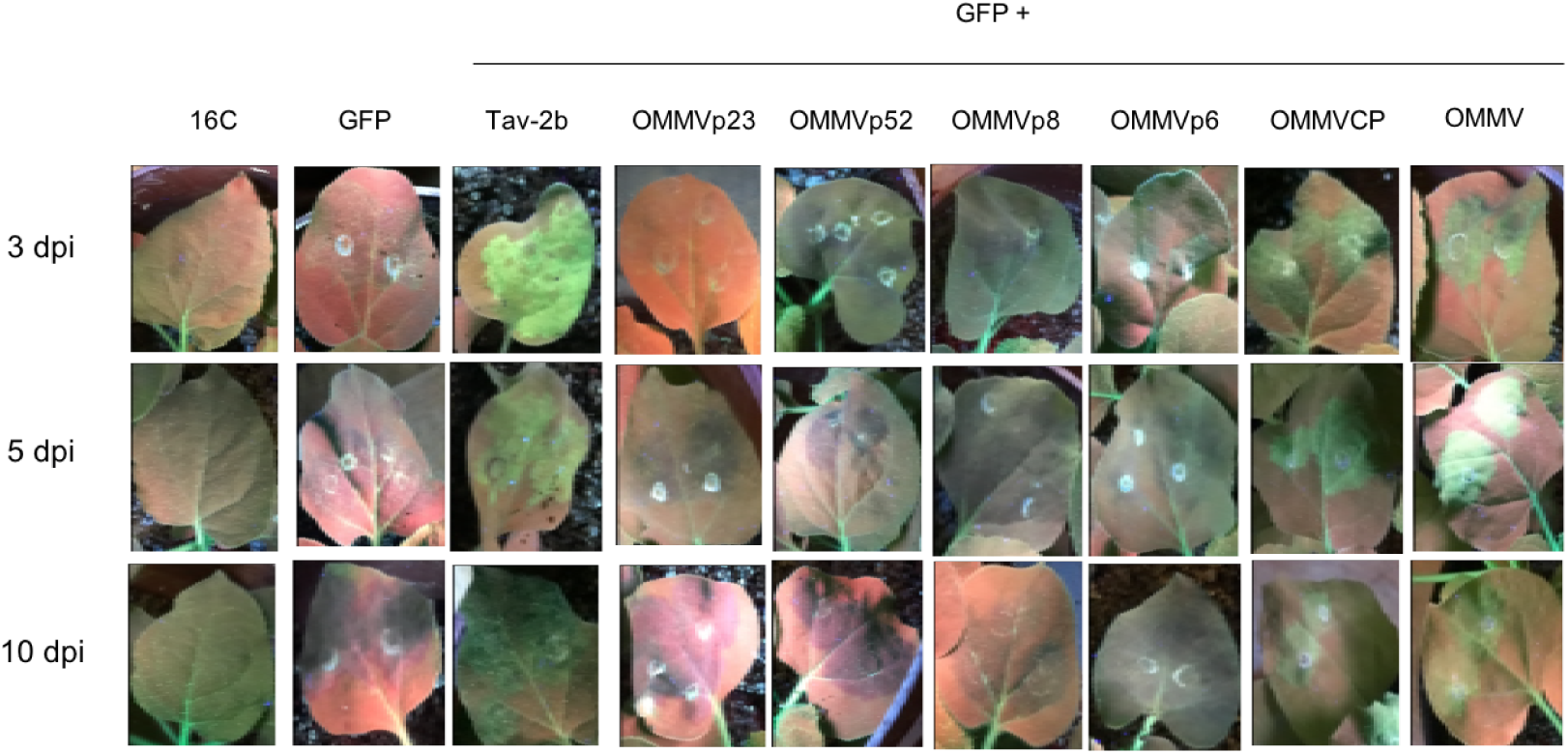
Visual observation of leaves from transgenic *N. benthamiana* 16C plants under UV light at 3, 5 and 10 days postinfiltration (dpi). Non infiltrated (16C), infiltrated with Agrobacterium tumefaciens harbouring pK-GFP alone (GFP) and co infiltrated with pK-GFP plus: Tav-2b, OMMVp23, OMMVp52, OMMVp8, OMMVp6, OMMVCP and OMMV.

In the positive control, leaves co-infiltrated with pk_GFP and the strong suppressor Tav-2b presented bright fluorescence at 3 dpi (Figure 1, Tav 3 dpi) and fluorescence was maintained for the next 7 days, although less intense (Figure 1, Tav-2b 10 dpi).

In the co expression of OMMVp23, OMMVp52 and OMMVp8, at 3, 5 and 10 dpi, no fluorescence was observed under UV light and silencing began at 5 dpi and was maintained at 10 dpi.

At 3, 5 and 10 dpi, OMMVp6, OMMVCP and OMMV were able to suppress GFP silencing, as seen by the green fluorescence under UV light.

OMMVCP and OMMV showed the most intense suppressor activity at 5 dpi, whereas OMMVp6 showed the most intense suppressor activity at 3 dpi. However, OMMVp6 green fluorescence showed lower intensity at all times when compared to OMMVCP and OMMV and was almost undetectable at 10 dpi.

GFP fluorescence in the presence of pk_OMMV was reproducibly stronger than with each of the viral genes and similar to the levels observed for Tav-2b. However, in contrast to Tav-2b, where higher levels were observed at 3 dpi, a brighter fluorescence was observed at 5 dpi.

Monitoring of upper non infiltrated leaves showed systemic silencing at 15 dpi in all samples, suggesting that OMMVp6 and OMMVCP possess suppressor local RNA silencing activity but cannot suppress systemic silencing.

RT-qPCR showed that all samples presented C_T_ values within the linear calibration curves. Two reference genes (PP2 and FBox) were used to normalize target genes expression. The amplification efficiency and correlation coefficient (R^2^) of their calibration curves were 107.25% and 0.9943 for PP2 and 99.62% and 0.9978 for FBox. As expected by visual observations, RT-qPCR at 3 dpi showed higher GFP mRNA levels on plants co inoculated with pk_GFP + Tav-2b, + OMMVp6, + OMMVCP and + OMMV as compared to single inoculated with pk_GFP and non-inoculated 16C leaves (Figure 2), reaching the highest levels with the strong suppressor Tav-2b. Plants co infiltrated with pK_GFP and OMMVp23, OMMVp52 and OMMVp6 showed GFP mRNA levels higher than non-infiltrated 16C plants and similar to those obtained with single pK_GFP infiltrations. At 5 dpi, RT-qPCR showed a great reduction in GFP mRNA levels of GFP alone or in the presence of OMMVp23, OMMVp52 and OMMVp8 and at 10 dpi GFP mRNA levels decreased to almost undetectable levels.

**Figure 2.**
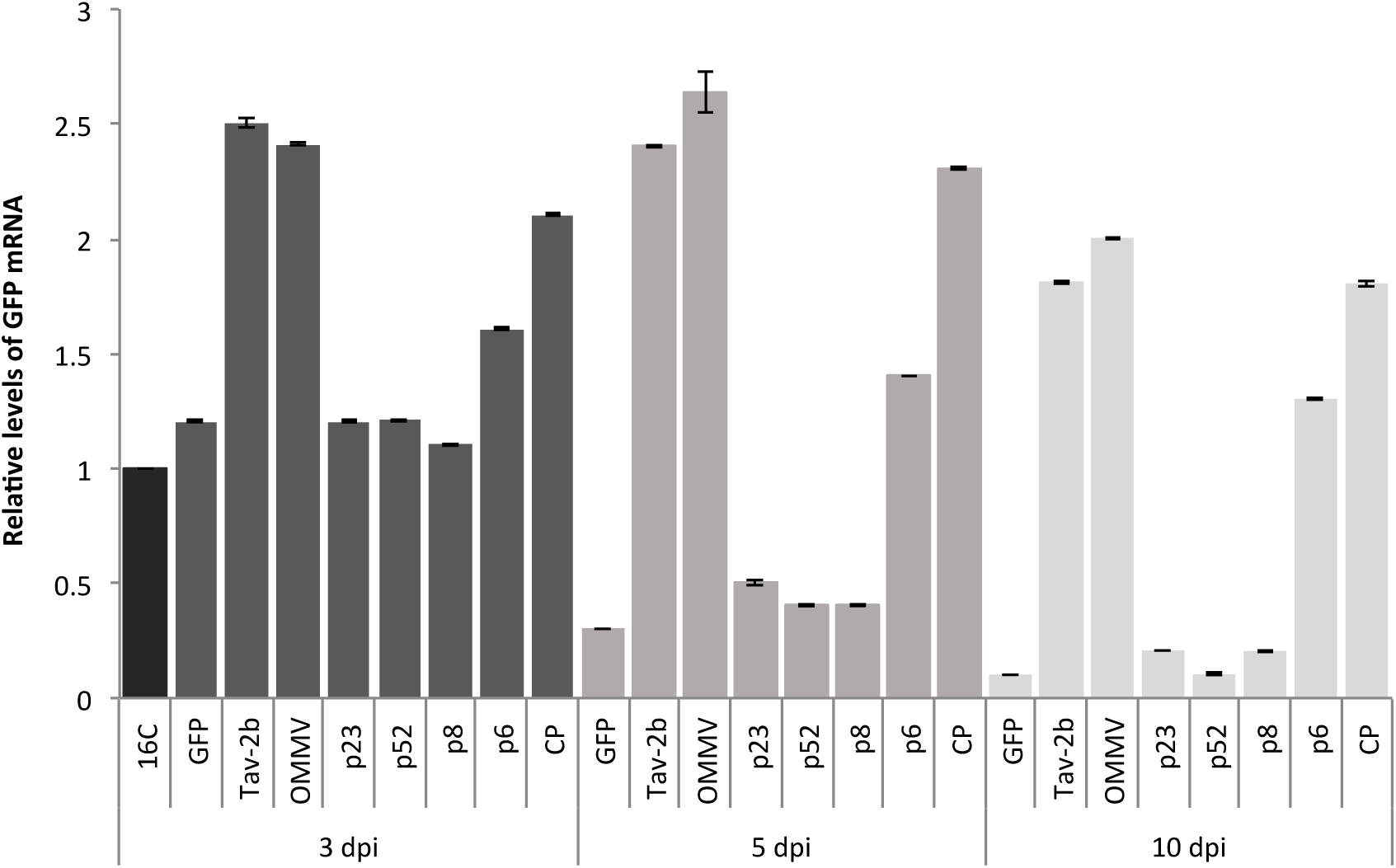
Mean relative levels of GFP mRNA ± standard error (SE) in infiltrated 16C *N. benthamiana* leaves at 3 dpi, 5 dpi and 10 dpi, determined by RT-qPCR and normalized by the levels of PP2 and F-box reference genes. 16C: 16C non-inoculated plants; GFP: single infiltration with pk_GFP; Tav-2b, OMMVp23, OMMVp52, OMMVp8, OMMVp6, OMMVCP and OMMV: co-infiltration of pk_GFP with the corresponding constructs. The GFP transgene level in a non-infiltrated *N. benthamiana* 16C plant was used as a standard and given a value of 1.

Consistent with visual observations, at 5 dpi, high GFP transcript levels were observed in patches expressing OMMVp6, OMMVCP, OMMV and Tav-2b, and at 10 dpi, all GFP levels decreased, nevertheless with higher levels than non-infiltrated 16C plants (from 0.3 in OMMVp6 to 1-fold greater in OMMV). GFP mRNA accumulation in OMMVp6 infiltrated patches was lower compared to OMMVCP and OMMV, suggesting that OMMVp6 is a weaker suppressor of silencing than OMMVCP.

The co infiltration of pk_GFP and OMMV reached the highest relative mRNA GFP levels (1.6-fold greater than non-infiltrated 16C at 5 dpi), and was higher than the levels obtained with the co infiltration with OMMVCP and OMMVp6 at all times tested, suggesting an enhanced suppressor activity resulting from the combined action of p6 and CP.

Plants co-infiltrated with pk_GFP and OMMVCP and pK_GFP and OMMV maintained the greenish patch for 11 and 14 days, respectively (data not shown).

At 3 and 5 dpi, expression in leaves co-infiltrated with pk_GFP and each OMMVp23, OMMVp52, OMMVp8 and OMMVp6, was similar to that of the strong silencing suppressor OMMVCP. This result confirms that proteins were being expressed and demonstrated that weak or no silencing suppression was not due to low protein expression.

PERMANOVA analysis revealed significant differences in the factors “ORFs” and “Time” (p < 0.0001). In addition, a significant interaction occurred between the two factors (“ORFs” and “Time”) (p < 0.0001). At 3 dpi the highest values (mean ±SE) of relative GFP mRNA abundance were 2.50 ± 0.01 in Tav-2b gene, followed by 2.41 ± 0.01 in OMMV, 2.10 ± 0.005 in CP and 1.60 ± 0.004 in p6 (Figure 2). Individual pairwise comparisons at 3 dpi revealed high variability of relative GFP mRNA abundance with significant differences between most ORFs (p < 0.05) (Table 3). Individual pairwise comparisons showed no significant differences between: *p* GFP *vs.* p23 > 0.9153, *p* p23 *vs.* p52 > 0.6178 and *p* p23 *vs.* p52 > 0.088.

At 5 dpi the highest values (mean ±SE) of relative GFP mRNA abundance were 2.64 ± 0.09 in OMMV followed by 2.40 ± 0.005 in Tav-2b, 2.30 ± 0.004 in CP and 1.40 ± 0.002 in p6 (Figure 2). Individual pairwise comparisons at 5 dpi revealed high variability of relative GFP mRNA abundance with significant differences between most of the ORFs (p < 0.05) (Table 3). Individual pairwise comparisons revealed no significant differences between: *p* Tav-2b *vs.* OMMV > 0.0518 and *p* p52 *vs.* p8 > 0.9977.

At 10 dpi the highest values (mean ±SE) of relative GFP mRNA abundance were 2.00 ± 0.005 in OMMV followed by 1.81 ± 0.005 in Tav-2b, 1.80 ± 0.01 in CP and 1.30 ± 0.003 in p6 (Figure 2). Individual pairwise comparisons at 5 dpi revealed high variability of relative GFP mRNA abundance with significant differences between most of the ORFs (p < 0.05) (Table 3). Individual pairwise comparisons revealed no significant differences between: *p* GFP *vs.* p52 > 0.3574, *p* p23 *vs.* p52 > 0.6807, *p* p23 *vs.* p8 > 0.6569, *p* p52 *vs.* p8 > 0.6799 and p Tav-2b vs. CP > 0.8368.

### OMMV resistance challenge

The presence of each construct was confirmed by RT-qPCR at 3dpi in plants infiltrated with Agrobacterium cultures carrying the constructs.

The appearance of disease symptoms was monitored at 2, 5, 10 and 16 dpi and recorded as a disease index scale. As shown in figure 3, at 2 dpi, plants expressing OMMV CP, OMMV p6 and both OMMV CP and p6 did not show any symptoms, whereas, 2 plants in negative control showed mild symptoms (DSI, 5%). At 5 dpi, plants from all groups showed symptoms, but only few plants of negative control group reached the disease index scale of 2, presenting the highest DSI (32.5%), followed by plants expressing p6 alone (22.5%). At 10 dpi and 16 dpi, control plants showed severe symptoms (DSI 75% and 100%, respectively). Plants expressing OMMV p6 showed a slight lower DSI at 10 dpi (60%) but also reached 100% at 16 dpi showing only a delay in the appearance of symptoms. At the same time points, plants expressing OMMV CP presented considerably lower DSIs (32.5% and 57.5%), indicating a tolerance to OMMV infection. In contrast, plants expressing both OMMV CP and p6 showed a maximum DSI of 20% at 16 dpi, indicating a very high tolerance to OMMV infection. Additionally, in the experiments, an average of 60% of the plants expressing OMMV CP and p6 did not present any viral symptoms at 16 dpi, suggesting a resistance to OMMV. Plants expressing OMMV CP also showed the existence of resistant plants, though in a lower level (20%). Plants were monitored until 30 dpi and the highest DSI was 60% in plants expressing OMMV CP and 30% in plants expressing both OMMV CP and p6, no plants reaching the maximum disease index.

**Figure 3.**
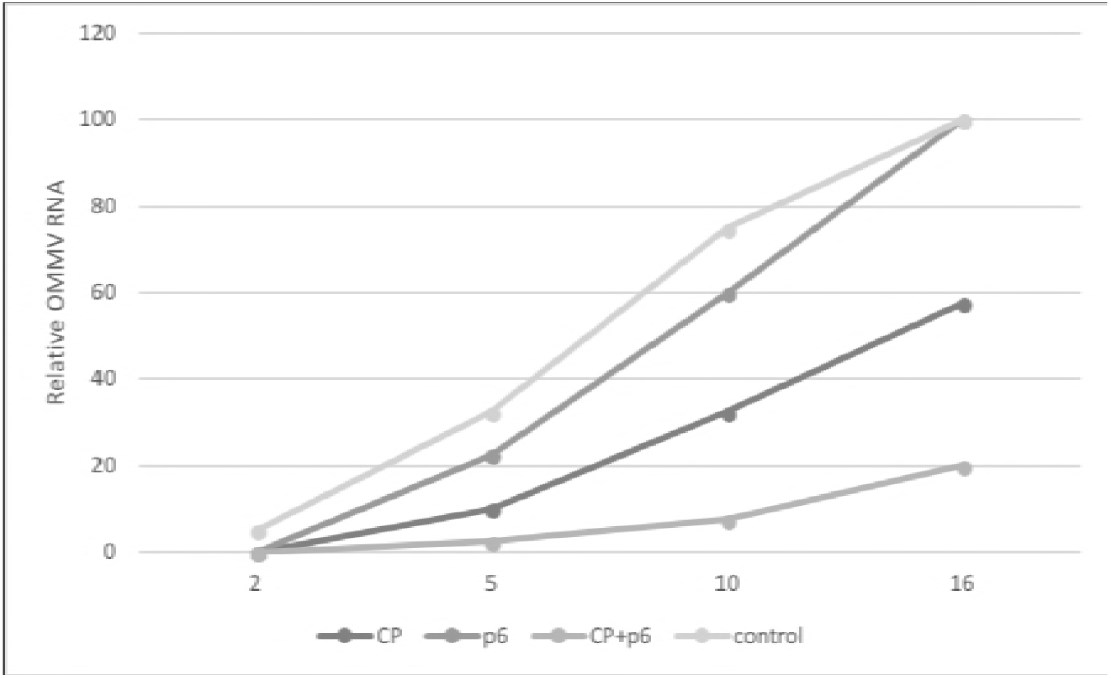
Disease severity of plants expressing OMMV CP, OMMV p6, OMMVCP + p6 and negative control at 2, 5, 10 and 16 days after inoculation of OMMV.

**Figure 4.**
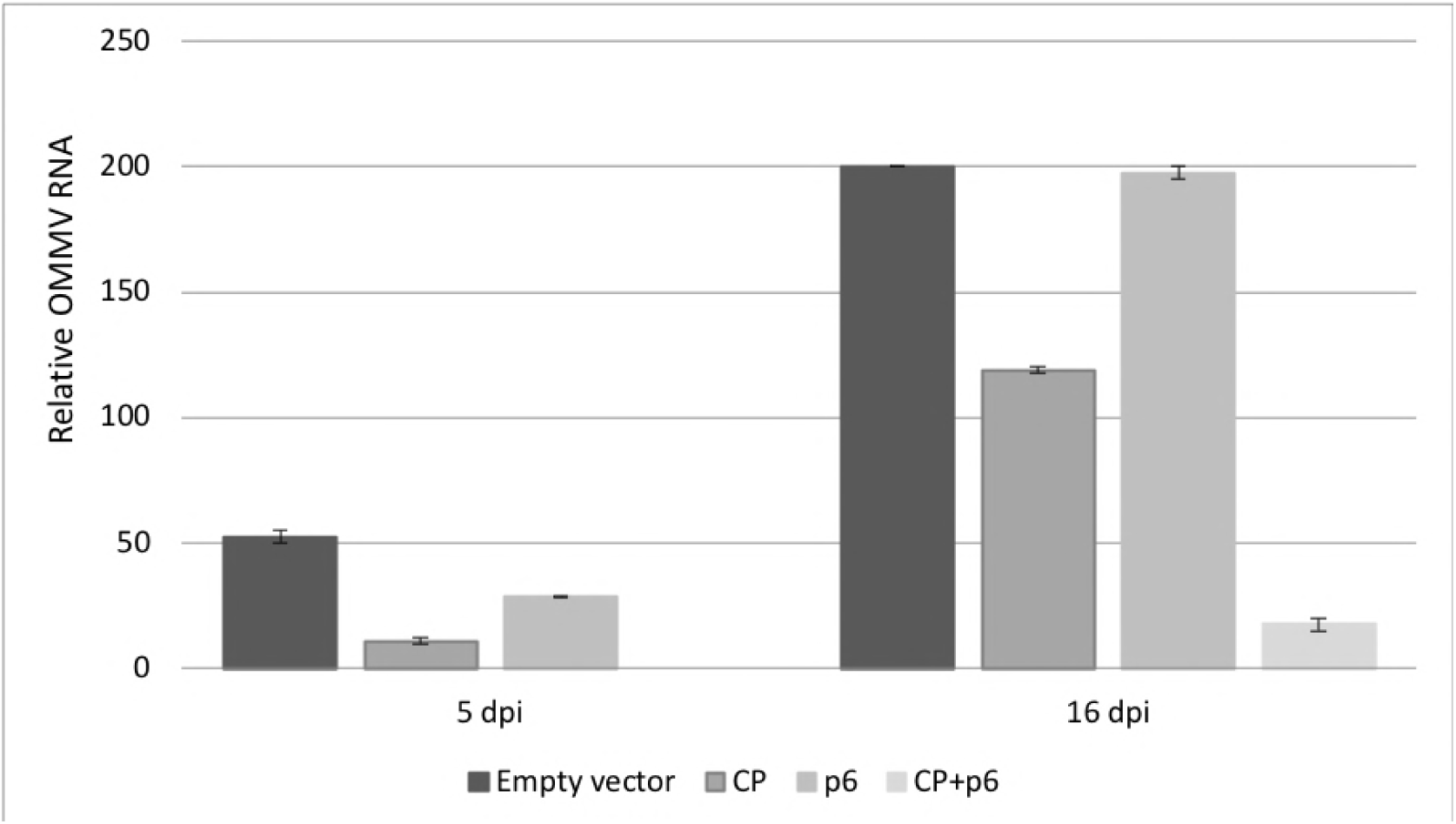
Viral accumulation levels in inoculated leaves at 5 and 16 days after OMMV inoculation.

Viral accumulation levels were determined by real time RT-qPCR and results are consistent with disease severity symptoms observed. Plants expressing OMMVCP and OMMVp6 individually resulted in significantly less virus accumulation in inoculated leaves at 5 dpi than when the empty vector was used. At 5 dpi no virus was detected in plants expressing both OMMV CP and p6. At 16 dpi, plants transformed with OMMV CP and plants transformed with both OMMV CP and p6, showed significant less OMMV accumulation than plants transformed with OMMV p6 and the negative control. Plants expressing both OMMV CP and p6 showed significant lower accumulation values than plants expressing OMMV CP. Although a slight lower viral accumulation was obtained in plants expressing OMMV p6 and empty vector at 16 dpi, differences were not significant.

Both visual observations and quantitative RT-PCR indicate that OMMV CP, and remarkably OMMV CP together with p6, attenuate viral symptoms and reduce viral accumulation levels.

## Discussion

Most plant viruses have evolved by encoding one or more silencing suppressors as a counter-response to the antiviral plant gene silencing defence mechanism (11). Members of the *Tombusviridae* family share common features, but, show different strategies to suppress silencing. Tombusvirus p19, the equivalent p14 of the aureusvirus, the replicase proteins of the dianthovirus and the CP of the carmoviruses (alpha-and beta-) have been identified as Tombusviridae silencing suppressors.

The high diversity of viral suppressors within *Tombusviridae* family did not anticipate a more likely suppressor for the alphanecrovirus OMMV. The closest related genera, the carmoviruses, use the CP as silencing suppressor, however, in opposition to the carmoviruses CP, the CP of necroviruses lacks a protruding domain and has low similarity to the CP of carmoviruses. In addition, the genome of OMMV does not present proteins equivalent to tombusvirus p19 and aureusvirus p14 and the necroviruses replicase has very low similarity to the replicase of the Dianthovirus (39).

In this study, the complete OMMV genome was screened for the presence of a gene with potential RNA silencing suppression activity. Coinfiltration assays using *N. benthamiana* 16C line, GFP-transformed, revealed that the full genome of OMMV has a suppressor activity similar to the strong suppressor Tav-2b. When each of the individual proteins were tested separately, CP and p6 were capable of inhibiting dsGFP-induced local silencing, at different levels, as seen by enhanced GFP fluorescence and GFP mRNA levels, suggesting a coordinate and complementary action of both CP and p6 as silencing suppressors that result in an enhanced activity when CP and p6 are co expressed. The analysis of each protein separately demonstrated that most suppressor activity is due to the CP, with p6 showing a much lower inhibition of local RNA silencing.

This finding is in line with previous data on OMMV mutants containing several changes in CP sequence, which exhibited different levels of symptoms in indicator plants, suggesting a clear role of that protein in symptom modulation (26).

The combination of several functions in the same protein may be advantageous for the virus. In particular, the silencing suppression function in the CP may have advantage since it is the viral genome most exposed to host defence machinery, due to the mode of virus replication (40).

This is the first time that two independent proteins were found to act as silencing suppressors in a member of Tombusviridae family similarly to that shown in some members of Closteroviridae (41) and Fimoviridae (42).

The existence of different viral suppressor proteins in one virus seems to be an advantage in targeting different pathways of host defence at cellular level as it allow for a more successful host infection. Viruses with single suppressors may only gain this advantage in mixed infections with others that possess distinct silencing suppressors with different targeting modes thus causing a more effective interference with the plant defence mechanism that result in increased symptoms, higher viral accumulation and favour cell to cell movement (43-45). This has been observed with PVX and CMV that encode distinct suppressors that target both intracellular and intercellular silencing leading to enhanced viral accumulation and symptoms characteristic of synergistic effect, in double infections (46).

Synergistic effects have been seen between OMMV and the close related OLV-1 during co-infection. OMMV is only acquired from the soil when OLV-1 is present, in which case both viruses are able to invade the plant and spread throughout the plant causing systemic symptoms instead of the typical local lesions induced by each virus separately (47). In fact, it is interesting to notice that both OMMV silencing suppressors are predicted to be involved in movement, p6 in cell to cell movement and CP in systemic movement (48).

The discovery of OMMV viral suppressors have prompted us to explore their use in the development of OMMV resistant plants through pathogen-derived resistance (PDR) based on RNA silencing.

The development of viral resistant plants has been a matter of study for many years and different levels of resistance have been obtained varying with the type of molecules and viral genome region used to trigger RNA silencing (49-51). As an alternative to transgenic plants and to overcome their associated biosafety and legislative constrains, transient RNA silencing systems have been developed consisting on the direct delivering of different RNA silencing molecules into plants (52-54).

The expression of self-complementary hairpin-RNA, in opposition to single sense or antisense constructs, induces a high level of RNA silencing in plants due to the production of dsRNA through the transcription of the hairpin structure; in addition, the presence of an intron in between the complementary regions stabilizes the construct and enhances silencing efficiency (55-57).

Tests were conducted to find out whether expression of artificial RNAs targeting the suppressors CP and p6 could counteract the suppressor function and confer resistance to OMMV using hairpin constructs. Two hairpin constructs targeting each the p6 and the CP were agroinfiltrated in plants to transiently express long pathogen related dsRNAs and preactivate plant RNA silencing machinery through the procession of dsRNAs into siRNAs responsible for the degradation of invading viral RNA.

For the construction of the p6 hairpin construct, the full p6 was used; for the CP hairpin construct, the first 141 nt of the CP were removed (OMMV CPminus141) in order to reduce the potential biosafety risks associated to the production of CP molecules as well as to exclude CP mediated resistance events, still a long fragment was used so that a higher number of potentially active virus-specific sRNAs would be present and specifically target the coding region of CP.

This study shows that the expression of OMMV CPminus141 and OMMV p6 interfered with the multiplication of OMMV as plants expressing them resulted in a highly substantial reduction in viral accumulation and symptoms attenuation, proving their effectiveness against OMMV infection. The percentage of resistant plants obtained here (60%) is similar to the highest percentages of viral resistance obtained in other studies (54, 57, 58).

Plants transiently expressing OMMV CPminus141 interfered with the multiplication of OMMV however it was not as effective against OMMV infection as plants expressing both that and OMMV p6 as just 20% of the plants were OMMV resistant. No resistant plants were obtained when OMMV p6 was being expressed however a slight delay in the appearance of symptoms occurred when compared to control. These results show that OMMV can counteract these mechanisms by inhibiting silencing. Only when plants expressed both OMMV suppressors, CP and p6, a high efficient RNA silencing mechanism was triggered. This has also been observed for CTV where resistance is only achieved when the three viral silencing suppressors were targeted simultaneously (59). Findings obtained here contribute to increase the knowledge on resistance induction frequencies and are of great importance for the future development of OMMV resistant plants, either transgenic or, in alternative, in transient systems by directly delivering RNA silencing molecules into plants, as to overcome the lack of regulatory framework and strong opposition associated to transgenic plants. Additionally, resistance to other close related viruses may be achieved as viruses with less than 15-20% differences at sequence level may confer protection to each other (60, 61). Such small differences exist between OMMV and other necroviruses such as TNV-D and OLV-1, whose co infections are frequent in nature (62). It is common for plants to be invaded by several viruses, and this study shows the most efficient OMMV transgenes that confer resistance to OMMV, which may be used in multiple combined sequences assembled with other viruses expected to infect the crop to achieve protection and a broader resistance against a wide range of viruses.

Besides helping to understand the interaction between viruses and hosts, the knowledge on viral suppressors may also have other commercial impacts as in the production of heterologous proteins using bioreactor plants. Host RNA silencing has shown to reduce the efficiency of gene expression in plants (63). In this way, the co-expression of a silencing suppressor and the target gene may be an attractive option to reduce problems in transgene expression. This is essential to respond to the continuous increase in the demand to produce large amounts of recombinant proteins for the industry for which plants are one of the most effective and safe systems for large scale production.

## Materials and Methods

### OMMV silencing suppressor genes

#### Generation of constructs

Plasmid DNA containing the viral full-length of an OMMV clone (27) was used for amplification of OMMV p23, p52, p8, p6 and CP ORFs as well as full-length OMMV. Each amplified sequence was cloned into pDONR™221 (Invitrogen) through a Gateway recombination reaction in accordance to manufacturer’s instructions. Primers used in the reactions are listed in table 1. Genes were transferred from pDONR221 to pK7WG2 binary vector (28), under the promotor CaMV 35S, through LR recombination. Confirmation of the correct sequences was done by sequencing the constructs. The 2b suppressor gene of *Tomato aspermy virus* (TAV), used as positive control, and green fluorescent protein (GFP) (m-gfp5-ER) (29), used as a silencing inducer, were cloned as described previously (30).

**Table 1:**
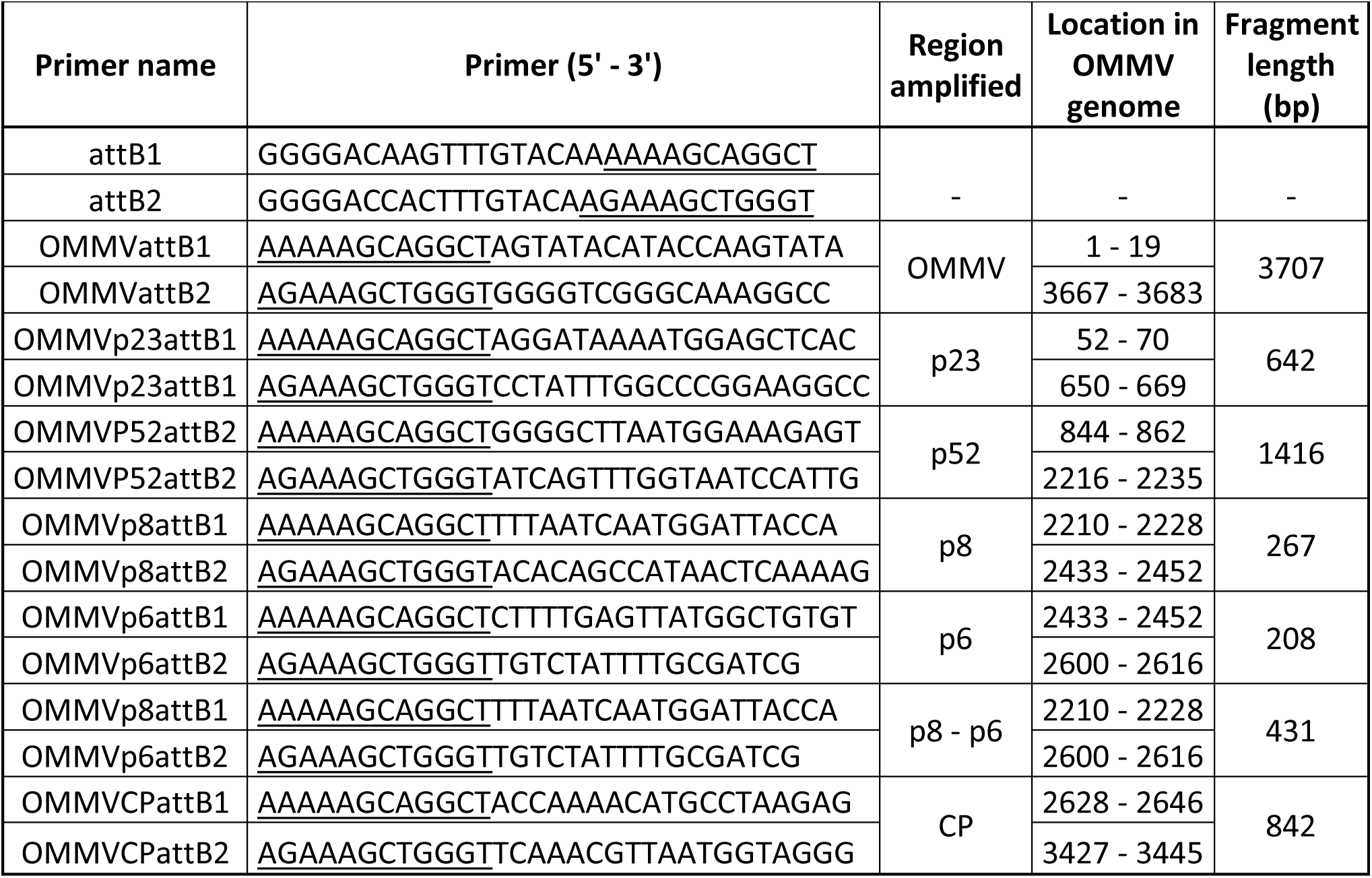
Primers used in the Gateway recombination reactions. Recombination sequences specific for the gateway system are underlined.

Binary vectors were transformed into competent *Agrobacterium tumefaciens* strain GV3101/C58C1, carrying pMP90 Ti plasmid which confers resistance to gentamycin. *A. tumefaciens* cultures were grown individually on Luria-Bertani (LB) medium supplemented with gentamycin, spectinomycin and rifampicin at 50 μg mL^-1^ each, 10mM MES and 20 μM acetosyringone at 28 °C, and 200 rpm until reaching an OD_600_ of 0.5. Cells were precipitated, re-suspended in 10 mM MgCl_2_ (pH 5.6), 10 mM MES and 100 μM acetosyringone, and kept in the absence of light at room temperature for 1 h before infiltration (31).

#### Agrobacterium co-infiltration assays and GFP imaging

Silencing suppressor assays were based on the previously described system (31). Briefly, when leaves of *N. benthamiana* 16C line plants carrying a copy of 35S:GFP:TNos transgene are agroinfiltrated with *A. tumefaciens* carrying the construct 35S GFP, the green fluorescent signal disappears under UV light due to GFP silencing. However, if the construct 35S GFP is co infiltrated with a silencing suppressor, the fluorescence does not disappear and may even become more intense due to the inhibition of gene silencing caused by the suppressor.

For transient expression assays, Agrobacterium cultures carrying each construct were infiltrated into leaves of 4-week-old *N. benthamiana* 16C line plants, gently provided by David Baulcombe (University of Cambridge, UK). Single and co-infiltrations assays were performed using a 5 mL needleless syringe. Single infiltration consisted on pK_GFP in a 15 mL suspension of *A. tumefaciens*. For co-infiltration assays, Agrobacterium cultures containing each construct individually, including Tav-2b, and GFP were mixed in 1:1 v/v ratio before agroinfiltration, centrifuged and ressuspended in a final volume of 15 mL. Three leaves per plant and ten plants per each construct were infiltrated. Plants were observed during 12 days post infiltration (dpi). Each experiment was repeated three times.

GFP fluorescence of infiltrated leaves and whole plants was examined using a long-wavelength UV lamp (Blak-Ray B-100AP, UVP) and photographed with a digital camera (Sony α100 DSLR-A100K).

#### RNA extraction and Real-Time RT-PCR

Total RNA was extracted from randomly selected agroinfiltrated leaves at 3, 5 and 10 dpi, using RNeasy Plant Mini Kit (Qiagen) in accordance with the manufacturer’s instructions. The quality and concentration of all RNA preparations were determined by Nanodrop 2000c (Thermo Scientific).

For reverse transcription, 1 μg of total RNA was used in a 20 μL reaction using Maxima^®^ First Strand cDNA Synthesis Kit for RT-qPCR (Thermo Scientific) in accordance to manufacturer’s instructions.

Primers were designed using Primer Express 3.0 software for Real-Time PCR (Applied Biosystems) using the default parameters for the software (Table 2). Quantitative assays of GFP mRNA were performed by real time RT-PCR (RT-qPCR), carried out on a 7500 Real Time PCR System (Applied Biosystems).

**Table 2:**
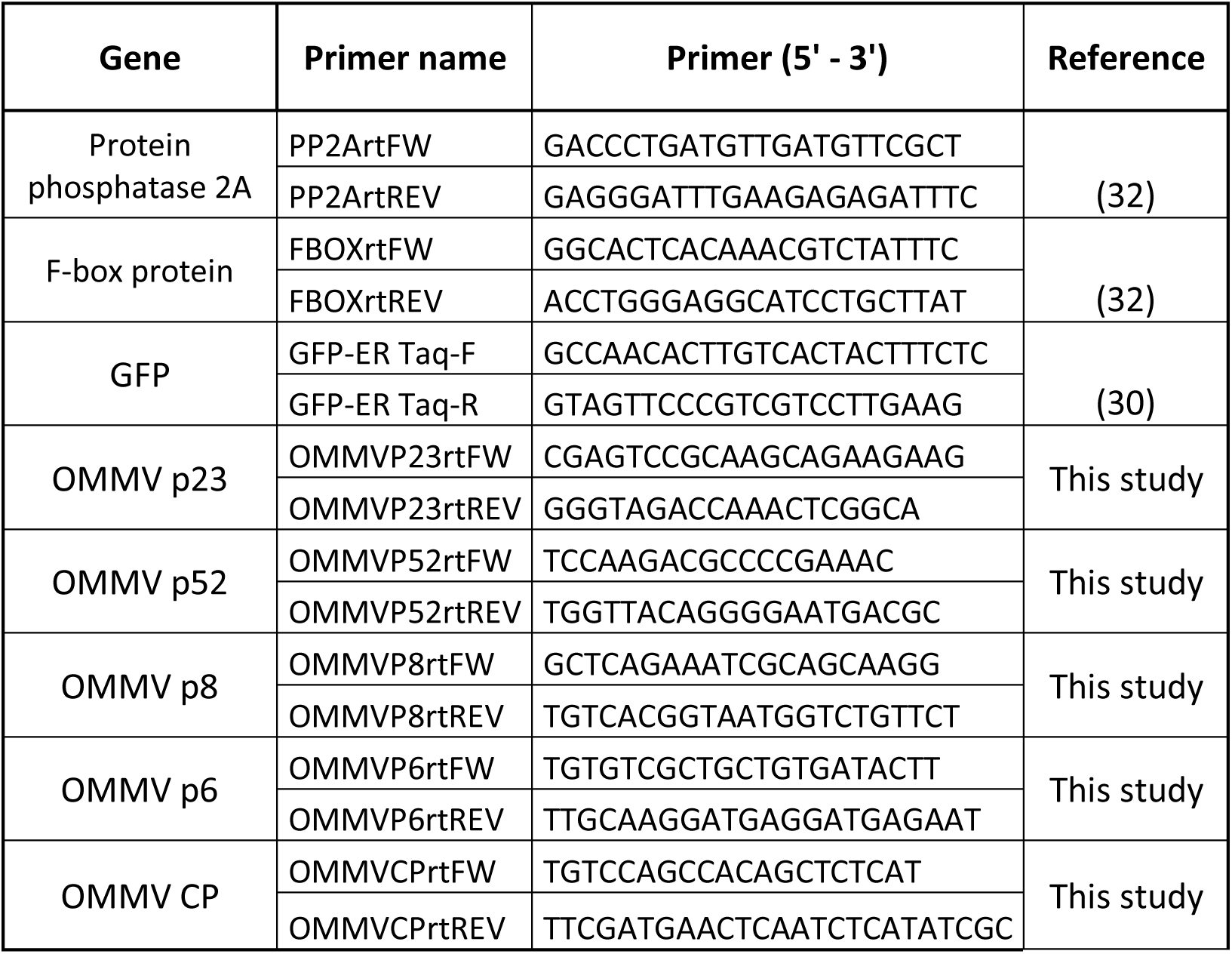
Primers used in RT-qPCR

**Table 3:** Details of the two-factor PERMANOVA Pair-Wise tests with “ORFs” (8 levels, fixed) and “Time” (3 levels, fixed) for all variables analysed. Bold values highlight significant effects (p < 0.05).

| Pair-Wise tests<br>ORFs | 3 dpi | 5 dpi | 10 dpi |
| --- | --- | --- | --- |
| GFP vs Tav-2b | **0.0001** | **0.0001** | **0.0001** |
| GFP vs OMMV | **0.0001** | **0.0001** | **0.0001** |
| GFP vs p23 | 0.9153 | **0.0001** | **0.0001** |
| GFP vs p52 | 0.6178 | **0.0001** | 0.3574 |
| GFP vs p8 | **0.0002** | **0.0001** | **0.0001** |
| GFP vs p6 | **0.0001** | **0.0001** | **0.0001** |
| GFP vs CP | **0.0001** | **0.0001** | **0.0001** |
| Tav-2b vs OMMV | **0.0129** | 0.0518 | **0.0001** |
| Tav-2b vs p23 | **0.0001** | **0.0001** | **0.0001** |
| Tav-2b vs p52 | **0.0001** | **0.0001** | **0.0001** |
| Tav-2b vs p8 | **0.0001** | **0.0001** | **0.0001** |
| Tav-2b vs P6 | **0.0001** | **0.0001** | **0.0001** |
| Tav-2b vs CP | **0.0001** | **0.0004** | 0.8368 |
| OMMV vs p23 | **0.0001** | **0.0001** | **0.0001** |
| OMMV vs p52 | **0.0001** | **0.0001** | **0.0001** |
| OMMV vs p8 | **0.0001** | **0.0001** | **0.0001** |
| OMMV vs p6 | **0.0001** | **0.0002** | **0.0001** |
| OMMV vs CP | **0.0001** | **0.0134** | **0.0001** |
| p23 vs p52 | 0.6137 | **0.0016** | 0.6807 |
| p23 vs p8 | **0.0001** | **0.0017** | 0.6569 |
| p23 vs p6 | **0.0001** | **0.0001** | **0.0001** |
| p23 vs CP | **0.0001** | **0.0001** | **0.0001** |
| p52 vs p8 | **0.0001** | 0.9977 | 0.6799 |
| p52 vs p6 | **0.0001** | **0.0001** | **0.0001** |
| p52 vs CP | **0.0001** | **0.0001** | **0.0001** |
| p8 vs p6 | **0.0001** | **0.0001** | **0.0001** |
| p8 vs CP | **0.0001** | **0.0001** | **0.0001** |
| p6 vs CP | **0.0001** | **0.0001** | **0.0001** |

A mRNA GFP 147-bp fragment, OMMV ORFs (p23, p52, p8, p6, CP) as well as full length OMMV, were amplified. The protein endogenous control genes phosphatase 2A (PP2) and F-box protein (F-box) were used as internal standards.

RT-qPCRs were carried out with 12.5 μL of 2 × SYBR Green PCR Master Mix, 0.3 μM of each primer and 12.5 ng of cDNA per sample, prepared in 96-well plates and run for 40 amplification cycles comprising a 15 s denaturation at 95 °C, followed by a 1 min at 60 °C step. A negative control with no template and three technical replicates were considered.

Cycle threshold (C_T_) values were determined using the fit-point method and the Applied Biosystems 7500 software with a fluorescence threshold arbitrarily set to 0.1. The relative level of GFP mRNA was determined using the amount of GFP mRNA from 16C non-inoculated plants as reference level. At the end of the qPCR, melt curve analysis was conducted to validate the specificity of the primers. A standard curve for each gene was constructed for relative expression level estimation and data was normalized by PP2 and F-box (reference genes).

To exclude the possibility of weak or no silencing suppression being due to low levels of respective protein expression in infiltrated leaves, quantitative assays of mRNA of each the proteins were performed by RT-qPCR as described previously for GFP.

#### Data analysis

Univariate and multivariate analyses were performed using the PRIMER v6 software (33) with the permutational analysis of variance (PERMANOVA) add-on package (34), to detect significant differences (p < 0.05) in the relative GFP mRNA abundance between; “ORFs” p23, p52, p8, p6, CP, OMMV, GFP and Tav-2b, and “Time” day 3, day 5 and day 10. A two-way PERMANOVA was applied to test the null hypotheses that no significant differences existed between “ORFs” and “Time”. PERMANOVA analyses were carried out with the following two-factor design, ORFs; p23, p52, p8, p6, CP, OMMV, GFP and Tav-2b (eight levels, fixed); and Time; day 3, day 5 and day 10 (three levels, fixed). The data were square-root transformed in order to scale down the importance of highly values of relative GFP mRNA abundance. The PERMANOVA analysis was conducted on a Bray-Curtis similarity matrix (35). If the number of permutations was lower than 150, the Monte Carlo permutation p was used. Whenever significant interaction effects were detected, these were examined using a posteriori pairwise comparisons, using 9999 permutations under a reduced model.

### OMMV resistance challenge

A full length cDNA of OMMV (36) was used for in vitro transcription using RiboMax™ Large Scale RNA Production System-T7 (Promega). Following transcription, DNA templates were removed by digestion with DNase and transcripts were purified by extraction with phenol:chloroform (5:1) acid equilibrated (pH 4.7) (Sigma) and ethanol precipitated.

Synthesized RNA was mechanically inoculated onto 6-8 leaf stage *N. benthamiana* plants, maintained in a growth chamber at 23 °C with a 16 h photoperiod, for viral propagation. OMMV infected leaves were then ground in cold 0.1 M, sodium phosphate buffer, pH 6.0 (1:3w/v), filtered, clarified in the presence of organic solvents, concentrated by differential centrifugation and further purified by ultracentrifugation through sucrose density gradients (37). The concentration of viral preparations was determined by Nanodrop 2000c (Thermo Scientific) prior to inoculation.

Hairpin constructs of CP and p6 were constructed based on the CP without the first 141 nt to exclude CP mediated resistance events (using OMMVCP-141attB1: 5’ <u>AAAAAGCAGGCT</u>ATCCTAGATCTTCTGGGCTAAGC and OMMVCPattB2) and on the full p6 (using OMMVp6attB1 and OMMVp6attB2), respectively. hpRNA-CP and hpRNA-p6 constructs were obtained through LR recombination from each pDONR221-CP and pDONR221-p6, as described previously, to pHELLSGATE12 (38), placed in sense and antisense directions to produce self-complementary dsRNAs. Confirmation of the correct sequences was done by sequencing of the constructs after linearization with *Cla*I which cleaves within the intron.

15 μg of purified OMMV were inoculated onto two fully expanded carborundum dusted leaves of 4-week-old *N. benthamiana* 16C line plants 3 days after infiltration with Agrobacterium cultures carrying pHELLSGATE12-CP and pHELLSGATE12-p6, as described for transient expression assays. Inoculated plants were grown in the conditions mentioned above. The presence of each construct was confirmed by RT-qPCR at 3dpi as shown previously.

Ten plants were infiltrated with pHELLSGATE12-CP, ten with pHELLSGATE12-p6, ten with both constructs and ten with the empty vector to be used as negative control. All 40 plants were mechanically inoculated with OMMV. Experiments were repeated two times.

Plants were monitored daily for 30 days for symptom development. A four-grade disease scale was adopted to describe OMMV symptoms along time: 0, no symptoms; 1, mild chlorotic mosaic; 2, intense chlorotic mosaic; 3, necrotic mosaic; 4, pronounced leaf necrosis and deformation. Disease severity was evaluated on 2 dpi, 5 dpi, 10 dpi and 16 dpi for each batch of plants infiltrated with the different constructs, as follows: Disease Severity Index (DSI) = (SUM of all disease ratings/(Total number of ratings*Maximum disease grade))*100.

Total RNAs were extracted from inoculated leaves from 3 randomly selected plants, at 5 dpi and 16 dpi. Virus accumulation was determined by quantitative real time PCR analysis using OMMVp23 primers (Table 2) as described above.

Univariate and multivariate analyses were performed as described above, using the PRIMER software to detect significant differences (p < 0.05) in the virus accumulation between; “ORFs” p6, CP and p6 + CP, and “Time” 5 dpi and 16 dpi. A two-way PERMANOVA was applied to test the null hypotheses that no significant differences existed between “ORFs” and “Time”. PERMANOVA analyses were carried out as previously, with the following two-factor design, ORFs; p6, CP and p6 + CP (three levels, fixed); and Time; 5 dpi and 16 dpi (two levels, fixed). The data were square-root transformed in order to scale down the importance of highly values of virus accumulation.

## Acknowledgements

This work was funded by National Funds through FCT-Foundation for Science and Technology under the Project UID/AGR/00115/2013 and by the European Union through the European Regional Development Fund, under the ALENTEJO 2020 (Regional Operational Program of the Alentejo) through the project Enhancing the Performance of Portuguese Olive Cultivars (OLEAVALOR) ALT20-03-0145-FEDER-000014. C. Varanda was supported by a post-doctoral fellowship (SFRH/BPD/76194/2011) from the Foundation for Science and Technology (FCT), funded by QREN – POPH – Typology 4.1 – co-funded by MES National Funding and The European Social Fund. The authors would like to thank CSIRO for providing pHELLSGATE vector.

The funders had no role in study design, data collection and interpretation, or the decision to submit the work for publication. The authors declare that there is no conflict of interest.

CV, GN and MRF designed the studies. CV, PM and MRF designed the experiments. PM performed the statistical analyses. MDC and MIC contributed to discussions. CV, PM, GN and MRF contributed to the interpretation of the results. CV and PM performed the figures and drafted the manuscript. All authors contributed for the writing of the final manuscript.

